# Variable Presence of an Evolutionarily New Brain Structure is Related to Trait Impulsivity

**DOI:** 10.1101/2024.10.23.619912

**Authors:** Ethan H. Willbrand, Samira A. Maboudian, Matthew V. Elliott, Gabby M. Kellerman, Sheri L. Johnson, Kevin S. Weiner

## Abstract

**Background:** Impulsivity is a multidimensional construct reflecting poor constraint over one’s behaviors. Clinical psychology research identifies separable impulsivity dimensions that are each unique transdiagnostic indicators for psychopathology. Yet, despite this apparent clinical importance, the shared and unique neuroanatomical correlates of these factors remain largely unknown. Concomitantly, neuroimaging research identifies variably present human brain structures implicated in cognition and disorder: the folds (sulci) of the cerebral cortex located in the latest developing and most evolutionarily expanded hominoid-specific association cortices.

**Methods:** We tethered these two fields to test whether variability in one such structure in anterior cingulate cortex (ACC)—the paracingulate sulcus (PCGS)—was related to individual differences in trait impulsivity. 120 adult participants with internalizing or externalizing psychopathology completed a magnetic resonance imaging scan and the Three-Factor Impulsivity Index. Using precision imaging techniques, we manually identified the PCGS, when present, and acquired quantitative folding metrics (PCGS length and ACC local gyrification index).

**Results:** Neuroanatomical-behavioral analyses revealed that participants with leftward or symmetrical PCGS patterns had greater severity of Lack of Follow Through (LFT)—which captures inattention and lack of perseverance—than those with rightward asymmetry. Neuroanatomical-functional analyses identified that the PCGS co-localized with a focal locus found in a neuroimaging meta-analysis on a feature underlying LFT. Both quantitative folding metrics did not relate to any impulsivity dimension.

**Conclusions:** This study advances understanding of the neuroanatomical correlates of impulsivity and establishes the notion that the topographical organization of distinct, hominoid-specific cortical expanses underlie separable impulsivity dimensions with robust, transdiagnostic implications for psychopathology.

## Introduction

Identifying the neuroanatomical correlates of psychopathology is a major goal of clinical and evolutionary neuroscience. Given the extensive evidence that comorbidity is the norm rather than the exception among individuals suffering from mental illness, research has sought to understand transdiagnostic risk variables shared across disorders (1). Impulsivity, generally speaking, is defined by trait-like tendencies toward poor constraint over one’s behaviors (2,3). A crucial advance in recent decades is the reproducible finding that rather than being a unidimensional construct, impulsivity is a set of separable, but overlapping dimensions (2,4,5) with robust, yet dissociable patterns of covariance with internalizing and externalizing psychopathology (3,6–8). Although there are multiple measurement approaches to impulsivity, thousands of studies now validate self-report measures that distinguish between impulsive responses to states of high emotion and impulsivity scales without reference to emotion. Impulsivity dimensions characterized by losses of self-control during elevated emotional states, such as the Feelings Trigger Actions (FTA) and Pervasive Influence of Feelings (PIF) dimensions, are often highlighted for their transdiagnostic associations with psychopathology (9–11). However, the Lack of Follow Through (LFT) dimension, which is characterized by distractibility and lack of perseverance to one’s goals (5), also explains unique variance in numerous externalizing and inattentive symptoms (6,12,13). Therefore, elucidating the shared and unique neuroanatomical underpinnings of FTA, PIF, and LFT is critical for improving the specificity of clinical interventions for these different forms of impulsivity.

Neurobiological investigations into impulsivity, broadly speaking, have consistently pointed to the prefrontal cortex (PFC) across species and developmental stages (14,15). In humans, although patients with PFC lesions may present with increased impulsivity (16–18), the neuroanatomical correlates of impulsivity remain quite heterogeneous within the relatively vast human PFC (19). Review articles did not identify reproducible neuroanatomical correlates within PFC for studies using traditional gray matter volume and cortical thickness measurements (20,21). However, recent studies indicate that other structural metrics may be better fit to unlock the neuroanatomical correlates of trait impulsivity (22). Our recent investigations identified that emotion-related impulsivity dimensions (FTA and PIF) were related to a regional-level quantitative measure of cortical folding (local gyrification) of the orbitofrontal cortex (OFC) (23). Furthermore, regional PFC surface area yielded stronger effects than cortical thickness did with both emotion-related and non-emotion-related impulsivity dimensions in the Adolescent Brain and Cognitive Development (ABCD) sample (*N* = 11,052) (24). These promising regional-level findings have set the stage to investigate whether specific PFC structures are linked to emotion-related and non-emotion-related forms of impulsivity.

Hominoid-specific structures, which include many of the folds (sulci) comprising the cerebral cortex, are appealing candidates, as they may serve as biomarkers for human-specific aspects of cognition—especially in brain regions such as PFC that have expanded substantially throughout evolution. Crucially, focusing on sulci circumvents contentions regarding analogous and homologous PFC areas across species for two main reasons (22). First, there are contentions regarding what criteria are ideal to define specific brain areas in association cortices. Second, while cerebral cortices vary substantially in size and folding across species, comparing brain areas across species using multiple criteria is useful to determine potential analogous and homologous areas despite these differences. Nevertheless, the parcellation of the cerebral cortex into areas can vary depending on the methods used. For example, Brodmann parcellated the cerebral cortex into 52 areas based on cytoarchitecture (25), while his mentors, the Vogts, parcellated the cortex into 180 areas based on myeloarchitecture (26). The Vogts included sulci, while Brodmann did not. Contrary to this variability and the potential moving boundaries of cortical areas based on the data used, sulci do not move [and a majority of the cerebral cortex is buried in sulci (27–30)]. Thus, sulci may serve as stable landmarks to measure both cross-sectionally and longitudinally across various age groups and clinical populations, and importantly, sulcal morphology and the presence or absence of sulci has been linked to different aspects of cognition in these groups (30–44).

Accordingly, the primary aim of this study is to precisely tease apart specific prefrontal structures that are also identifiable in non-human hominoids and that map onto distinct dimensions of impulsivity in humans. One method that is well-matched to achieve this goal is the use of high-resolution cortical reconstructions generated from T1-weighted MRI images to identify and quantify features of cortical sulci (39,44–46). Indeed, we recently found that one of the emotion-related impulsivity dimensions (FTA) was related to the depth of specific OFC sulci bilaterally (47) and other groups identified that variations in the sulcal organization of OFC are altered across multiple disorders (34,48). Here, we extend this previous work by considering variations in the sulcal organization of anterior cingulate cortex (ACC) and impulsivity. We targeted ACC given its key roles in emotion-cognition integration (49), of relevance for FTA and PIF, and error monitoring and attentional control (50–52), which have been shown to be impaired in the components of impulsivity measured in LFT: lack of perseverance and attention deficits (5,53).

In terms of sulcal organization, ACC is marked by the consistent cingulate sulcus (CGS) and the variably-present paracingulate sulcus (PCGS) (54–56)—an evolutionarily new [great ape specific (57,58)] tertiary [emerging in the second/third trimester (59–61)] sulcus above the CGS (**Figure 1A**). In neurotypical populations, the PCGS is present in at least one hemisphere in approximately 60-70% of participants and shows leftward asymmetry - being more common in the left than right hemisphere (33,36,54–57). The PCGS also shows more leftward asymmetry in males than females (55,56,62–64). In addition, inter-hemispheric asymmetry in PCGS presence is related to executive function and can be altered in disorder (31,33,36,65–73). For example, in healthy populations, an asymmetric PCGS pattern has been associated with better inhibitory control efficiency (31,65–67). In clinical populations, findings can be summarized in the following four ways. First, there is increased rightward asymmetry and overall less asymmetry in patients with schizophrenia (68–71). Second, patients with obsessive-compulsive disorder are less likely to have a left hemisphere PCGS (72). Third, patients with bipolar disorder are less likely to have a PCGS in both hemispheres (73). Fourth, PCGS length is reduced and predictive of hallucinations in patients with schizophrenia (32,74). However, it is unknown if features of the PCGS are trait markers for transdiagnostic features of psychopathology.

**Figure 1.**
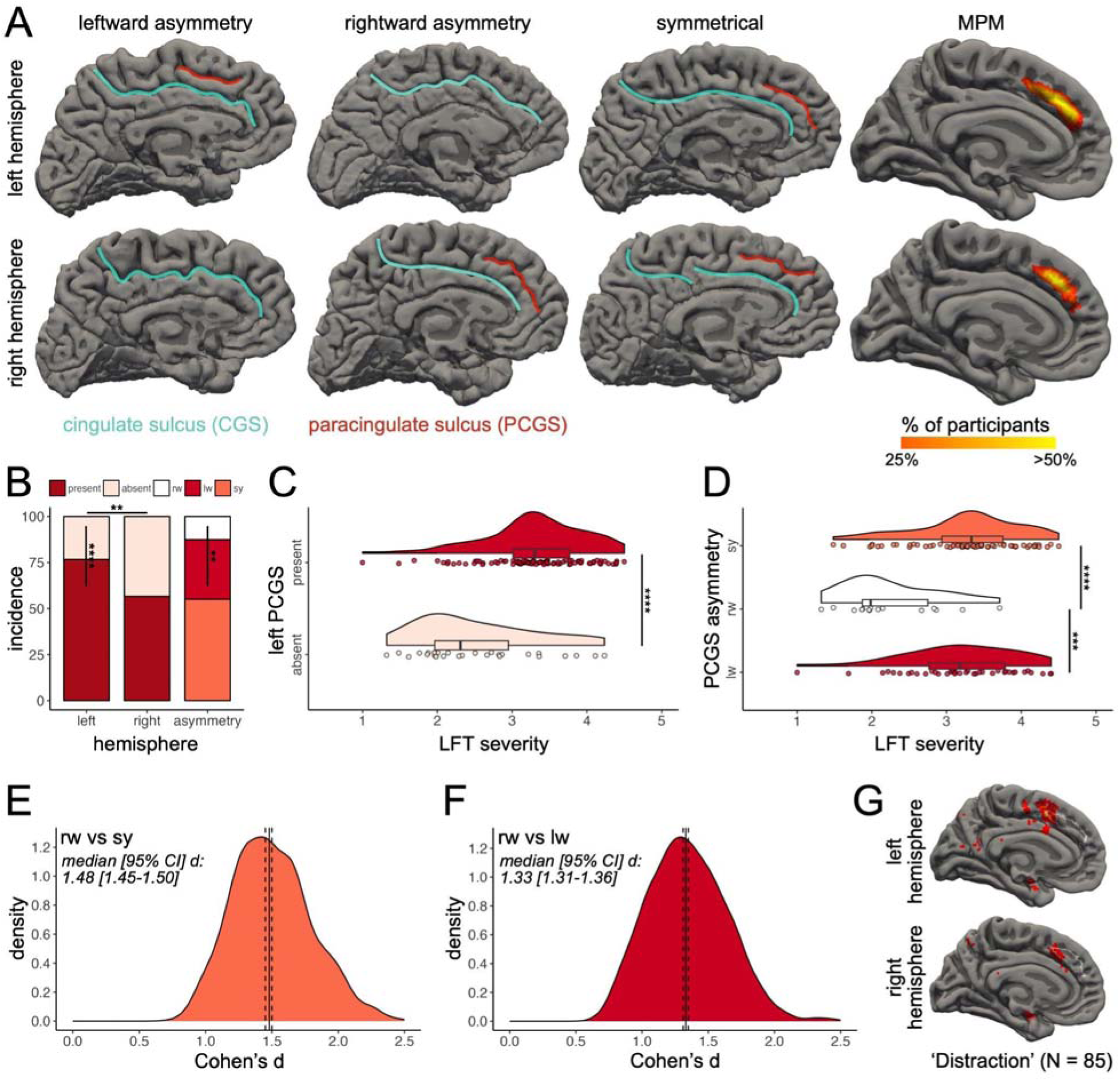
The paracingulate sulcus is related to non-emotional impulsivity. **(A)** Hemisphere presence/absence (rows) and asymmetry (columns) of the PCGS in three example participants in both hemispheres on the wrinkled (pial) cortical surface reconstructions (CGS: blue; PCGS: red). The rightmost column shows the transdiagnostic maximum probability map (MPM) of the PCGS in both hemispheres (thresholded at 25% overlap for visualization purposes). **(B)** Bar plots showing PCGS incidence (present/absent) and asymmetry rates. **(C)** Raincloud plot (118) showing LFT severity as a function of left PCGS presence. **(D)** Same as (C), but for PCGS asymmetry. **(E-F)** Distribution of the iteratively sampled effect size (Cohen’s d) of the significant effects in (D) with the median (black line) and 95% CI (dotted lines). **(G)** Pial fsaverage cortical surfaces displaying overlap of a whole-brain FDR-corrected (*p* =□0.01) uniformity-test meta-analysis z-score map of the “distraction” term (N = 85 studies; heatmap) and PCGS MPMs (white outline). Abbreviations are as follows: Lack of Follow Through (LFT); leftward (lw); rightward (rw); symmetrical (sy). Asterisks indicate the following *p* value thresholds: \*\**p* < .01, \*\*\**p* < .001, \*\*\*\**p* < .0001.

To this end, we first documented the intra- and inter-hemispheric presence of the PCGS in a transdiagnostic adult sample (N = 120, ages 18-55; **Table 1**) with varying severity of internalizing and externalizing syndromes, studied in our prior work (23,47). We then explored the relationship between three impulsivity factors and variability in PCGS presence. Finally, we assessed whether quantitative features of the PCGS (length) and ACC (local gyrification index) were linked to each impulsivity factor.

**Table 1.** Participant Characteristics (n = 120)

| Characteristic | n (%) or Mean (SD) [Range] |
| --- | --- |
| Gender |  |
| Female | 79 (65.8%) |
| Male | 36 (30%) |
| Nonbinary | 5 (4.2%) |
| Race |  |
| Asian/Asian American | 37 (30.8%) |
| Native Hawaiian/Pacific Islander | 1 (0.9%) |
| Black/African American | 8 (6.7%) |
| White/European American | 52 (43.3%) |
| Other/Multiple Races | 16 (13.3%) |
| Declined to Answer | 6 (5%) |
| Ethnicity |  |
| Hispanic or Latinx | 23 (19.2%) |
| Not Hispanic or Latinx | 97 (80.8%) |
| Age, Years | 28.3 (8.6) [18-55] |
| Education, Years | 15.7 (2.3) [12-21] |
| SCID-5 Lifetime Diagnosis |  |
| Major depressive disorder | 97 (80.8%) |
| Anxiety disorder | 81 (67.5%) |
| Alcohol use disorder | 25 (20.8%) |
| Substance use disorder | 24 (20%) |
| More than one disorder | 82 (68.3%) |
| Impulsivity Subtype |  |
| Pervasive influence of feelings | 3.68 (0.76) [1.92-5] |
| Feelings trigger action | 2.81 (0.74) [1.14-4.52] |
| Lack of follow through | 3.11 (0.78) [1.00-4.50] |

*Note.* Abbreviations are as follows: Structured Clinical Interview for DSM-5 (SCID-5).

## Methods and Materials

### Participants

The present study consisted of 120 adults (18-55 years old; 66% Female) who participated in a parent study that was approved by the UC Berkeley Committee for the Protection of Human Subjects. Full participant details are documented in **Table 1** and the **Supplementary Materials**.

### Behavioral Data Acquisition

Trait impulsivity scores were calculated using the well-validated Three Factor Impulsivity Index. Factor analyses consistently support three impulsivity factors with strong internal consistency for each scale (5,6,13,75,76). The first factor, FTA, reflects the tendency toward rash action or speech while experiencing overwhelming positively or negatively valenced emotions. FTA consists of three subscales: the Negative Urgency scale (2), Positive Urgency Measure (77), and Reflexive Reaction to Feelings scale (5). The second factor, PIF, is derived from three scales measuring cognitive and motivational responses to mostly negative emotion: the Generalization (78), Sadness Paralysis (5), and Emotions Color Worldview (5) scales. LFT measures two forms of impulsivity without reference to emotions: the Lack of Perseverance (2) and Distractibility (5) scales. Each factor is measured on a Likert scale of one to five, with higher scores indicating higher severity. The descriptive statistics of the impulsivity factors are included in **Table 1**. All impulsivity factors were unrelated to participant age and gender (**Table S1**).

### Imaging Data Acquisition

Participants were scanned using a 3T Siemens TIM Trio MRI scanner (Siemens Healthineers) at the University of California, Berkeley Brain Imaging Center. Sagittal T1-weighted structural images were acquired using a 32-channel receiver head coil and a 6.1-minute magnetization-prepared rapid gradient-echo sequence. Scanning parameters are as follows: repetition time = 1900 ms, echo time = 2.89 ms, field of view = 256 mm, voxel size = 1-mm^3^ isotropic voxels, and parallel acquisition technique mode = GRAPPA, with acceleration factor PE = 2. Cortical surface reconstructions were generated from the T1-weighted images using FreeSurfer (v6.0) (79–81). Manual PCGS identification was performed on cortical surface reconstructions in each individual (46,82–84).

### Neuroanatomical Data Acquisition

#### Paracingulate Sulcus (PCGS) Identification and Classification Criteria

The PCGS is a variably-present sulcus located dorsal and parallel to the CGS (**Figure 1A**). The PCGS was identified and classified according to its intra- and inter-hemispheric presence/absence following established criteria (32,38,65,66,68,71,85–88) that is documented in the **Supplementary Materials**. See **Figure 1A** for examples of intra- and inter-hemispheric PCGS presence/absence. We also generated probability maps for the PCGS in this transdiagnostic sample (**Supplementary Materials**; **Figure 1A**).

#### Relating the Paracingulate Sulcus to a Meta-Analysis of fMRI Studies Associated with LFT

Although fMRI research has yet to examine the functional correlates of LFT, to situate the PCGS within putative functional representations of LFT to guide future research, we leveraged the Neurosynth meta-analysis platform (89). We first searched the database for terms associated with LFT [e.g., perseverance (2) and distractibility (5)]. The only term identified was ‘distraction.’ Next, we downloaded the uniformity test meta-analysis maps for the search term ‘distraction’ (N = 85 studies). The uniformity test map was generated from a χ2 test comparing voxel activation in the studies containing the term to the expected activation if it were uniformly distributed across the gray matter. Finally, these maps were projected to fsaverage surface space with the *mri_vol2surf* FreeSurfer function so the PCGS probability maps could be spatially related.

### Quantification and Statistical Analysis

All statistical tests were implemented in R (v4.1.2) (90). We briefly overview the analyses here; full statistical details are in the **Supplementary Materials**.

#### Qualitative Neuroanatomical Analyses

We began by detailing PCGS incidence against multiple features as this has yet to be documented in a transdiagnostic sample (to our knowledge). We specifically examined whether PCGS incidence differed as a function of (i) hemisphere (within and between), (ii) Structured Clinical Interview for DSM-5 (SCID-5) Lifetime Diagnosis, (iii) age, and (iv) gender. We then tested whether each impulsivity index differed as a function of (i) intra-hemispheric PCGS presence and (ii) inter-hemispheric asymmetries in PCGS presence.

#### Quantitative Neuroanatomical Analyses

We implemented two parallel analyses to examine the relationship between impulsivity and quantitative features of the PCGS (length, in mm) and ACC subregions (local gyrification index). Given that all results were null, the full methods and statistical analyses, as well as the data and results for these two analyses, are included in the **Supplementary Materials** (including **Tables S2–S7** and **Figures S1–S3**).

## Results

### A Significant Leftward Bias in Paracingulate Sulcus Presence in a Transdiagnostic Sample

Overall, at least one PCGS was present in 66% of participants in at least one hemisphere (**Figure 1B**). The incidence of the PCGS significantly differed between hemispheres (76% of left and 56% of right hemispheres contained a PCGS; χ2(1)□=□9.92, *p* = .001, 2-sample test for equality of proportions with continuity correction; **Figure 1B**). Within hemisphere, the PCGS was significantly more present than absent in the left hemisphere (χ2(1) =□33.08, *p =* .000000008, 1-sample proportions test with continuity correction), but not in the right hemisphere (χ2(1)□=□1.88, *p* = .17, 1-sample proportions test with continuity correction). As is common in the field (31,68,71,91,92), we also quantified the inter-hemispheric asymmetry in PCGS presence (**Figure 1A and 1B**). There was a significant leftward bias in PCGS presence in this transdiagnostic sample (χ2(1) = 9.80, *p* = .001, McNemar’s test with continuity correction; **Figure 1B**). It has been observed that PCGS asymmetry is reduced in some disorders, but not others (36,68–73,93). As such, the presence of asymmetry observed here could be due to the wide range of syndrome variability in this transdiagnostic sample (**Table 1**). However, χ2 tests (with Yates’ continuity correction) identified that intra- and inter-hemispheric PCGS presence did not vary as a function of any of the SCID-5 Lifetime Diagnoses (**Tables S8–S11**). Multiple tests were also performed to evaluate the effects of age and gender on intra- and inter-hemispheric PCGS presence (**Supplementary Materials**). There were no significant effects (**Tables S12** and **S13**). To aid PCGS identification in non-neurotypical samples, we provide “transdiagnostic” probabilistic maps with the publication of this paper (**Data and code availability**; **Figure 1A**).

### The Paracingulate Sulcus is Associated with Non-Emotional Impulsivity

We then sought to relate the presence of the PCGS in each hemisphere separately to each impulsivity index (FTA, PIF, and LFT). An ANOVA with factors of left and right PCGS presence revealed a main effect of left PCGS presence for LFT scores (F(1, 117) = 23.87, *p* = .00001, η_p_^2^ = 0.17), in which the presence of the left PCGS was associated with an increase in LFT severity (**Figure 1C**). Given the unequal sample sizes between groups (**Figure 1C**), we conducted permutation testing, which confirmed the result (*p*_n=1,000_*= .0009). The fact that there were no other significant effects of PCGS presence on any other impulsivity index tested (**Table S14**) and the difference in AIC (**Supplementary Materials**) between the LFT model for PCGS presence and PIF and FTA models (ΔAIC_PIF-LFT_ = 13.80; ΔAIC_FTA-LFT_ = 7.45) indicates that the predictive value of PCGS presence is strongest for LFT.

Next, given the previously documented relationship between inter-hemispheric PCGS asymmetry and cognitive functioning and disorder (31,33,36,68–71), we ran a second ANOVA to probe the relationship of PCGS asymmetry (symmetrical, leftward asymmetry, and rightward asymmetry) to LFT scores, which revealed a main effect of PCGS asymmetry (F(2, 117) = 12.69, *p* = .00003, η_p_^2^ = 0.18; **Figure 1D**), that was confirmed by permutation testing (*p*_n=1,000_* = .0009). Post hoc pairwise comparisons showed that participants with a rightward asymmetric PCGS asymmetry had lower LFT severity than symmetric (*p* = .000002, d = 1.47) and leftward asymmetric (*p* = .0001, d = 1.28) patterns, with no significant differences between symmetric and leftward asymmetric patterns (*p* = .59; **Figure 1D**). To account for the impact of differences in sample size between the three PCGS asymmetry groups on the post hoc effect size tests, we iteratively sampled a size-matched subset of the symmetric and leftward asymmetric groups to the rightward sample 1,000 times, which confirmed the effect on behavior (**Figure 1E and F**). No other impulsivity index tested was related to PCGS asymmetry (**Table S15**) and the difference in AIC (**Supplementary Materials**) between the LFT model for PCGS asymmetry and PIF and FTA models (ΔAIC_PIF-LFT_ = 15.63; ΔAIC_FTA-LFT_ = 9.13) indicates that the predictive value of PCGS asymmetry is strongest for LFT. Finally, to link the PCGS to LFT functionally, a meta-analysis of neuroimaging research on the term ‘distraction’ (89), a concept related to LFT (5), putatively co-localizes with the PCGS (**Figure 1G**).

## Discussion

Trait impulsivity has shown robust relationships with both internalizing (e.g., depression) and externalizing (e.g., alcohol/substance use) disorders (3,6–8,13). The present findings extend the current picture of the neuroanatomical underpinnings of impulsivity in two key ways. First, they show a regional neuroanatomical dissociation between the non-emotional and emotional impulsivity constructs. Specifically, our prior work (in a largely overlapping sample) identified that OFC folding was related to the severity of emotion-related impulsivity (FTA and PIF) (23,47), whereas the present work identified that non-emotion-related impulsivity (LFT) was related to ACC folding. In combination, these findings highlight that the separability of impulsivity facets in psychological assessment is supported by dissociable neurobiological correlates in the folding of different cortical expanses. This dissociation may be a consequence of these regions being members of different cortical networks—for example, those implicated in hot versus cold executive function (94). Second, both studies found that rightward asymmetry in folding was associated with reduced impulsivity severity for their respective index, indicating that rightward asymmetry is likely a general neuroanatomical principle protecting against impulsivity—a hypothesis which can be tested in future research. Integrating these points together, future research can seek to examine the sulcal organization of different regions [e.g., lateral and ventromedial PFC (37,39,44,46,57,95)] to further fill in the neuroanatomical map of impulsivity and determine if these dissociations (emotional versus non-emotional impulsivity) and consistencies (rightward asymmetry associated with lower impulsivity severity) hold.

In contrast to our prior work in OFC (23), we identified null relationships between ACC LGI and all impulsivity dimensions. These results indicate that the observed behavioral effects are driven by the neurobiological processes underlying variations of specific sulci, and not regional measures of folding (like LGI). Complementing our recent follow-up study examining the morphology of specific OFC sulci to impulsivity (47), these results support the notion that these specific impulsivity dimensions are associated with different scales of folding. For example, PIF is related to OFC LGI, but not necessarily the morphology of specific OFC sulci, whereas FTA shows the opposite pattern in OFC (23,47). Future investigations should seek to identify if different neurobiological substrates underlie these different measures of cortical folding to provide further insight into the neuroanatomical-behavioral differences observed in this triad of recent studies.

The observation that variability in PCGS presence, but not length, was associated with LFT provides insight into when the neuroanatomical correlates of LFT likely develop. PCGS presence is a cortical feature that forms in the second to third gestational trimesters (59,60,96), is predominantly determined by in utero environmental factors (33), and the presence/absence of the PCGS does not change after birth (67,97). This is in contrast to quantitative features of sulci (e.g., depth, length, width, and surface area), which change with age (63,98–100). These neurodevelopmental differences indicate that, to an extent, features of the fetal environment (in contrast to the postnatal environment) likely play a major role in establishing this neuroanatomical-behavioral effect (36). Longitudinal studies are necessary to further explore the differential impact of these two environments on the presently documented (and to be uncovered) neuroanatomical correlates of LFT.

Translationally, these results extend the growing literature supporting relationships between PCGS presence and multiple psychiatric disorders [e.g., schizophrenia (68–71), obsessive-compulsive disorder (72), and bipolar disorder (73)] by showing that this cortical feature is also associated with LFT, a transdiagnostic predictor of psychopathology, not just the disorders themselves. Indeed, we observed direct effects for LFT, but not for psychiatric diagnoses, on PCGS presence (**Tables S8–S11**), indicating that these cortical features might be more sensitive indicators of this preclinical trait than diagnoses were. This is key given that LFT has been tied to diagnoses and real world preclinical outcomes, such as engaging in risky behavior (101). One possibility is that PCGS presence provides an early indicator of a continuum of impulsivity risk, which then might be expressed as psychiatric diagnoses in the context of other risk variables. This idea will require further testing given that statistical power was limited by the imbalanced distributions of those with and without a diagnosis (**Table 1**).

These findings also hold relevance to understanding the etiology of specific disorders. For example, these findings extend a functional neuroimaging meta-analysis indicating ADHD-related hypoactivation only in the right paracingulate cortex during attention tasks (102). Since the current findings show that rightward paracingulate asymmetry is protective against severe LFT, and prior research indicates that (i) LFT is strongly associated with inattentive symptoms in ADHD (12) and (ii) PCGS presence/absence alters ACC functional activity (103–105), future investigations can tether these threads to assess whether the PCGS is functionally and behaviorally implicated in ADHD severity.

In another vein, these results build upon studies in neurotypical samples demonstrating a link between PCGS presence and cognitive performance by showing that these relationships extend to transdiagnostic samples. These prior studies identified that asymmetry in PCGS presence (either asymmetry in general, or in some cases leftward asymmetry) was associated with better performance on multiple cognitive measures [e.g., inhibitory control (31,65–67), fluid intelligence (33), reality monitoring (106), and verbal fluency (63)]. In the present study, we found that rightward asymmetry was associated with lower LFT severity. The anatomical and functional differences in the brains of individuals with different asymmetries in PCGS presence are not well understood (36), highlighting the need for future research to elucidate the mechanisms underlying these observed relationships.

Of course, the reader may also be asking: How could sulcal patterning mechanistically relate to impulsivity? A potential answer lies in the empirical and theorized link between sulci and underlying cortical anatomy and function [reviewed in (22,36,107)]. It has been proposed that PCGS presence could reflect strengthened or otherwise altered local connectivity within paralimbic cortex (BA 32) and neighboring regions (BA 6, 8, and 9), with implications for vulnerability to various disorders relating to ACC function (38,68,71,86). Indeed, prior research in neurotypical samples has documented that PCGS presence relates to changes in the local cytoarchitectonic organization of gray matter (54,108), structural and functional connectivity (109,110), and brain function (103–105). In disease, it has been suggested that right PCGS presence may be protective against disease-related neurodegeneration in behavior-variant frontotemporal dementia by altering local connectivity patterns, leading to a delay in disease onset (38,86). Similarly, studies of PCGS presence in schizophrenia have proposed that lower left PCGS sulcation observed in the disorder may relate to weaker ACC connectivity; increased left PCGS sulcation may thus be protective (68,71). We propose a similar mechanism at play, in which local connectivity patterns associated with rightward PCGS asymmetry may be protective—that is, be related to lower LFT. However, direct investigation of this hypothesis is warranted, and it is important to note that structural and functional network alterations related to PCGS presence are likely only one of many features playing a role in impulsivity.

Although the neuroanatomical precision of the present study is an undeniable strength, it is not without its limitations. Specifically, the time-intensive nature and neuroanatomical expertise needed to manually identify sulci leads to two key limitations. First, the sample size is often limited (46,67,68,83,95,111), necessitating follow-up confirmatory analyses. Although automated methods are being developed to address this limitation, these methods still fall short of manual identification in accuracy (112–115). For example, current automated methodologies identify the PCGS with 70-80% accuracy (112,113). Therefore, the current best approach is one integrating multiple methods—such as manual definition guided by probabilistic maps and automated labels. Second, manually defining sulci often limits studies to one region or one sulcus, limiting the observation of more complex interactions between sulci within and between regions on behavior (36). Accordingly, it is necessary to investigate the neuroanatomical correlates of impulsivity in other cortical expanses where the sulcal organization is cognitively and functionally relevant [e.g., lateral PFC (39,44,67), ventromedial PFC (57,95), and lateral parietal cortex (84,111,116)]. Finally, the functional relationship observed in **Figure 1G** is limited by two features. The first is the inherent limitation of relating two group-level probabilistic locations together, as this may not fully represent the individual-level relationship (117). The second is that local folding, cytoarchitecture, and functional features of ACC are all impacted by PCGS variability (54,104,105,108–110). As such, we suggest that future research examining the functional correlates of LFT should i) be done at both the individual and group levels, ii) consider the location of the PCGS as a landmark for functional activity when present, and iii) consider how functional activity might change with PCGS variability.

In closing, the present findings highlight that PCGS patterning is a crucial cortical feature that should be considered in future studies to examine how multiscale anatomical and functional features give rise to psychopathology.

## Acknowledgments and disclosures

All authors report no biomedical financial interest or potential conflicts of interest. This work was supported by: NIH R01 MH110447 (SLJ), Brain and Behavior Research Foundation NARSAD 30738 (KSW), NSF CAREER 2042251 (KSW), and NIH MSTP T32 GM140935 (EHW). The funding agencies did not have a role in the study design, data collection and analysis, decision to publish, or preparation of the manuscript. We thank K. Timpano, K. Modavi, A. Dev, M. Robison, J. Mostajabi, S. Esmail, and B. Weinberg for their help with recruitment and data collection. We also thank N. Angelides, H.Y. Tsai, M. Andrews, and J. Giffin for their help with magnetic resonance imaging data acquisition.

## Author contributions

EHW, SAM, and KSW designed research; EHW, SAM, GMK, and KSW performed research; MVE and SLJ contributed neuroimaging and behavioral data; EHW, SAM, MVE, SLJ, and KSW wrote the paper; all authors gave final approval to the paper before submission.

## Data and code availability

All data and original code used for the present project will be publicly available on GitHub upon publication (https://github.com/cnl-berkeley/stable_projects). Any additional information required to reanalyze the data reported in this paper is available from the corresponding author (Kevin Weiner) upon request.

