## Supplementary Materials for "Variable Presence of an Evolutionarily New Brain Structure is Related to Trait Impulsivity"

### Additional Participant Details

Individuals were recruited from the general community who were experiencing a wide range of mental health symptoms, indicated by scoring greater than 5 on the Sheehan Disability Scale in regard to mental health (1). Individuals were excluded who had current alcohol or substance use disorders or a history of bipolar disorder or primary psychosis [as assessed by the Structured Clinical Interview for DSM-5 (SCID-5)], or daily use of marijuana or sedating medications, as well as those with lifetime head trauma resulting in loss of consciousness for five or more minutes, diminished cognitive abilities [Orientation Memory Concentration Test score of less than 7 (2)], MRI safety contraindications, neurological disorders, or inability to complete cognitive measures due to intellectual or language impairment. After participants completed informed consent procedures and urine toxicology screening, they completed diagnostic, behavioral, and neuroimaging sessions.

### Neuroanatomical Data Acquisition

**Paracingulate Sulcus Identification and Classification Criteria.** The assessment of ACC sulcal organization focused on the presence or absence of a variable tertiary sulcus: the paracingulate sulcus (PCGS). The PCGS was defined as the sulcus located dorsal to the CGS with a course parallel to the cingulate sulcus (CGS) (3–7). Here, we followed established criteria by Yücel *et al.* (7). As outlined by Yücel *et al.* (7), to mitigate ambiguity arising from the convergence of the PCGS and the CGS with the superior rostral sulcus, the anterior limit of the PCGS was identified as the point where the sulcus extends posteriorly from an imaginary vertical line perpendicular to the line connecting the anterior and posterior commissures (AC–PC) and parallel to the anterior commissure. Following prior work (3,8–16), a binary classification was employed to categorize the presence of PCGS in each individual hemisphere (right and left). The PCGS was designated as "present" if its length was ≥ 20 mm, and "absent" when no clearly developed horizontal sulcus elements parallel to the cingulate sulcus were observed reaching that minimum length. The PCGS depth also needed to be > 4 mm to be present. In the present study, we did not adopt the “present” (20 mm ≤ length < 40 mm) versus “prominent” (≥ 40 mm) distinction (5,7,17,18) given that PCGS length changes with age and disorder (12,19–23). The PCGS was independently identified by trained raters EHW and SAM (Cohen’s κ ± SE = 0.910 ± 0.031) and confirmed by a trained neuroanatomist (KSW; there were no disagreements on any sulci). For additional analyses, authors EHW and GKM also manually defined the PCGS as a .label file in each individual hemisphere using FreeSurfer's *tksurfer* tools. See **Figure 1A** for examples of binary PCGS presence/absence.

Measurement of length (in mm) was calculated by determining the longest geodesic distance along the cortical surface between any pair of vertices located on the boundary of the label, as in our prior work (24–26). In instances where sulci were identified as comprising multiple disconnected pieces on the cortical surface, the sulcal length was defined as the cumulative length of each individual sulcal component, excluding any annectant gyral component(s). Geodesic distance calculations on the fiducial surface were executed using algorithms implemented in the pycortex python package (https://gallantlab.github.io/) (27). The descriptive statistics of PCGS length in both hemispheres are as follows: left hemisphere (mean ± std = 43.00 ± 19.50 mm; range = 20.58-110.57 mm), right hemisphere (mean ± std = 42.40 ± 20.70 mm; range = 20.75-118.69 mm). PCGS length distributions in both hemispheres are depicted in **Figure S1A**.

Measurement of sulcal depth (in mm) was calculated from the sulcal fundus to the smoothed outer pial surface using a custom-modified version of an algorithm building on the FreeSurfer pipeline (28). The descriptive statistics of PCGS depth in both hemispheres are as follows: left hemisphere (mean ± std = 9.66 ± 2.14 mm; range = 4.66-14.10 mm), right hemisphere (mean ± std = 9.01 ± 2.71 mm; range = 4.19-15.10 mm). PCGS depth distributions in both hemispheres are depicted in **Figure S1B**.

Given the relevance of anatomical asymmetries to psychopathology (11,13,29–32), we also classified inter-hemispheric asymmetries in PCGS presence according to a standard nomenclature (11,13,18,32,33). The PCGS was asymmetrical if the PCGS definition in one hemisphere differed from the other. For example, if the PCGS was “present” in the left hemisphere but “absent” in the right, a participant would be classified leftward asymmetric (and vice versa for rightward asymmetric). A participant had a symmetric patterning if the PCGS classification in each hemisphere was the same (present and present or absent and absent). Examples of PCGS asymmetry are depicted in **Figure 1A**.

**Sulcal Probability Map Generation.** Leveraging our previously published pipeline (34), sulcal probability maps were calculated to display the vertices with the highest alignment across participants with the PCGS. PCGS label files were first transformed from the individual to the *fsaverage* surface. Once transformed to this common template space, the proportion of participants for whom a given vertex was labeled as the PCGS was then calculated. Unthresholded maps and constrained versions of these maps—that is, maximum probability maps (MPMs), which improve map interpretability (34)—are available (**Data and code availability**). The maps in **Figure 1** were thresholded at 25% for visualization purposes.

**Extracting Anterior Cingulate Cortex Gyrification Values.** We began by mapping three automatically-defined labels from the Desikan-Killiany-Tourville atlas corresponding to the location of the ACC broadly and the PCGS specifically (35), the superiorfrontal, caudalanteriorcingulate, and rostralanteriorcingulate, onto each hemisphere of each participant’s cortical reconstruction in FreeSurfer (**Figure S2A**). After, mean local gyrification index (LGI) values were extracted with the *mris_anatomical_stats* FreeSurfer function with the *pial_lgi* flag from each label in each hemisphere (36,37). LGI is a metric comparing the amount of cortex buried within sulci to the amount of cortex on the outer visible cortex, or hull (**Figure S2B**) (37). The LGI distributions in each region are visualized in **Figure S3**.

**Additional Quantification and Statistical Analysis**

All statistical tests were implemented in R (v4.1.2) (38). In each set of analyses, $p < 0.05$ was considered significant after controlling for multiple comparisons via the false discovery rate (FDR) method.

**Qualitative Neuroanatomical Analyses.** In the first set of analyses, we began by detailing PCGS incidence against multiple features as this has yet to be documented in a transdiagnostic sample (to our knowledge). 2-sample proportions tests (with continuity correction) were used to test for hemispheric differences in PCGS incidence and 1-sample proportions tests (with continuity correction) were used to report within-hemisphere incidence probabilities, as in prior work (39). McNemar's test for symmetry (with continuity correction) was used to test whether PCGS asymmetry in one direction was counterbalanced by equal cases with an asymmetry in the other, as in prior work (7). To assess potential drivers of variations in PCGS presence and asymmetry, chi-squared ($\chi 2$) tests (with Yates' continuity correction) were implemented for each SCID-5 Lifetime Diagnosis (**Table 1**). In addition, given prior work showing differences in PCGS presence and asymmetry on age and sex (5,7,17,19,40), we tested these relationships in our transdiagnostic sample. Two ANOVAs were implemented to determine whether i) hemispheric PCGS presence [factors: left PCGS presence (present/absent) and right PCGS presence (present/absent)] and ii) PCGS asymmetry [factor: PCGS asymmetry (leftward asymmetry, rightward asymmetry, symmetrical)] varied with age. $\chi 2$ tests (with Yates' continuity correction) were again implemented to evaluate the relationship between gender and PCGS presence and asymmetry.

We then ran analysis of variance (ANOVA) tests to compare the relationship between i) hemispheric PCGS presence [factors: left PCGS presence (present/absent) and right PCGS presence (present/absent)] and ii) PCGS asymmetry [factor: PCGS asymmetry (leftward asymmetry, rightward asymmetry, symmetrical)] and each impulsivity index. ANOVA effect sizes are reported as partial eta-squared ($\eta_p^2$). In all analyses, we submitted significant statistical results to random permutation testing. Permutation testing is a robust non-parametric alternative to standard parametric significance tests because it does not rely on distributional assumptions such as normality, homogeneity of variance, or random sampling. It is a powerful tool for evaluating test statistics from the modest sample sizes characteristic of neuroimaging studies. Using 1,000 permutations of the dataset, in which behavioral scores were randomly shuffled, we generated empirical null distributions of F-statistic, which we compared with the observed values from the selected model. Permutation testing yields exact *p*-values ($p^*_{n=1,000}$). Post hoc pairwise comparisons were conducted for any significant effects, and post hoc effect sizes are reported as Cohen's d (d). We assessed how the LFT models performed relative to the PIF and FTA models by measuring the difference of their Akaike Information Criterion (AIC) (41). When comparing two models, a ΔAIC greater than 2 indicates an interpretable difference between models, while a ΔAIC greater than 10 indicates a substantial difference, with the lower AIC value indicating the preferred model (42,43). Finally, in line with our prior work on sulcal patterning (44,45), to further determine whether the observed relationship between PCGS asymmetry and LFT was impacted by differences in sample size between participants with leftward (n = 39) and symmetrical (n = 66) patterns compared to rightward (n = 15), we iteratively sampled a subset of participants from the symmetrical and leftward asymmetry groups to match that of the rightward asymmetry ($n_{permutations}$ = 1,000). To evaluate the effect size, we report the median and 95% confidence interval for the effect size (Cohen's d).

Even though gender was unrelated to each impulsivity index and PCGS variability (**Tables S1** and **S13**), given the documented relationship between gender and PCGS presence and asymmetry (5,7,17,19,40), we also ran exploratory analyses to explicitly test whether the relationship between LFT and the left PCGS presence and asymmetry was impacted by gender. To do so, we included an interaction term for gender in the two aforementioned ANOVAs. Since this resulted in a null interaction for both left PCGS presence ($F(1,116) = 0.15$, $p = .69$, $\eta_p^2 < 0.01$) and asymmetry ($F(2,114) = 0.82$, $p = .44$, $\eta_p^2 = 0.01$), we do not discuss this beyond here.

**Quantitative Neuroanatomical Analyses.** To extend prior work showing that PCGS length is implicated in disorder [for example, in schizophrenia (12,23)], for the first set of quantitative analyses, we built two multivariate regression models to relate PCGS length to each impulsivity dimension. In the first model, the length of the left and right PCGS was included as predictor variables. The second model tested whether asymmetry in PCGS length of the left or right hemisphere predicted any of the impulsivity dimensions. To do so, we calculated the laterality ratio for each subregion in each participant as defined by Hill *et al*. (29). The laterality ratio was calculated with the following equation:

$$\frac{(Right-Left)}{(Right+Left)}$$

Positive values indicated longer length in the right hemisphere relative to the left hemisphere. Altogether, in the second model, the PCGS length laterality ratio was included as the predictor variable. Age and gender were not included in either model as they were unrelated to length (**Table S2**).

To mirror our prior work in OFC (30), for the second set of quantitative analyses, we built two similarly-structured multivariate regression models to test whether ACC LGI predicted any of the impulsivity dimensions. In the first model, the LGI of each ACC subregion was included as predictor variables. The right and left hemisphere LGI values for each subregion were included as separate variables in this model. The second model tested whether asymmetry in the LGI of the left or right hemisphere predicted any of the impulsivity dimensions via the laterality ratio

(29). Altogether, in the second model, the LGI laterality ratio of each ACC subregion were included as predictor variables. Age and gender were not included in either model as they were unrelated to LGI in all ACC subregions (**Table S3**).

### Supplementary Results

#### Impulsivity Dimensions Are Not Associated with Paracingulate Sulcus Length

Given that PCGS length is implicated in psychopathology severity (12,23), we also conducted adjunct analyses testing whether PCGS length was a predictor of impulsivity severity. Linear regressions with predictors of left and right PCGS length for each impulsivity dimension identified null results on all dimensions (**Table S4**). As in prior work (30), and to mirror the PCGS asymmetry analyses (**Figure 1D**), we also implemented the laterality ratio to assess the effect of hemispheric asymmetries in PCGS length on impulsivity (**Supplementary Methods**). Hemispheric asymmetry in PCGS length did not relate to any impulsivity dimensions (**Table S5**).

#### Impulsivity Dimensions Are Not Associated with Regional Anterior Cingulate Cortex Gyrification

To complement our prior work in OFC (30), we implemented a similar approach by extracting the mean LGI values of three subregions comprising ACC in the Desikan-Killany-Tourville atlas (**Figure S2**) (35). Constructing linear regressions relating the LGI values of each region in each hemisphere to each impulsivity dimension identified null results for all dimensions (**Table S6**). As in prior work (30), we calculated the laterality ratio (**Supplementary Methods**) to assess the effect of hemispheric asymmetry in ACC LGI on ERI. Linear regressions relating the laterality ratio of each region's LGI to each impulsivity dimension also showed null results for all dimensions (**Table S7**).

### Supplementary Figures

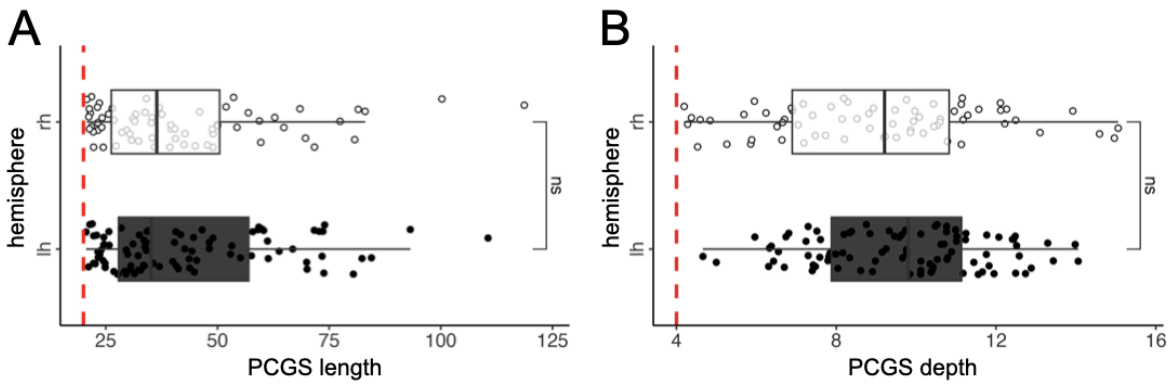


**Figure S1. Paracingulate length and depth in a transdiagnostic sample. (A)** Boxplot and scatterplot showing PCGS length (in mm; x-axis) as a function of hemisphere (y-axis). Dashed red line represents the standard 20 mm cutoff for identifying a PCGS as "present" (5,7,39). The significance of an effect of hemisphere on length (via a t-test) is shown. In contrast to neurotypical participants (19), we did not observe a significantly longer PCGS in the lh compared to rh. **(B)** Same as (A) but for PCGS depth (in mm). Dashed red line represents the standard 4 mm cutoff for identifying a PCGS as "present" (5,7,39). Abbreviations are as follows: paracingulate sulcus (PCGS), left hemisphere (lh), right hemisphere (rh). ns $p > .05$.

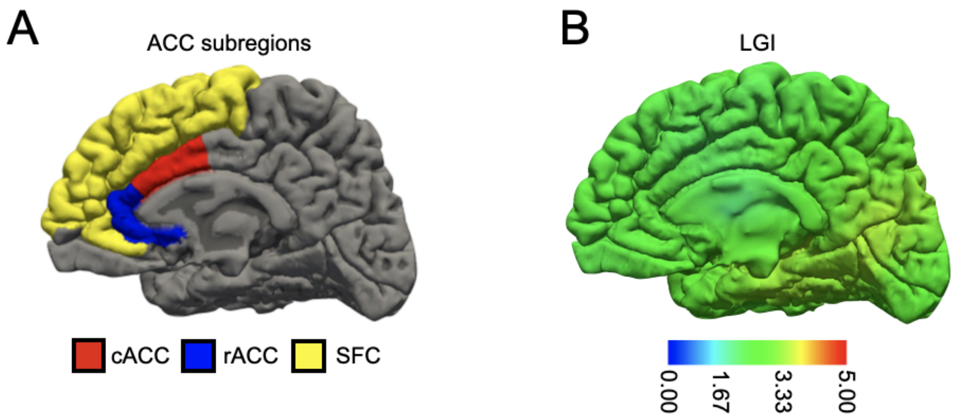


**Figure S2. Anterior cingulate cortex subregions and metric for quantitative neuroanatomical analyses. (A)** Three automatically-defined ACC subregions (colored according to the key) from the Desikan-Killany-Tourville atlas (35) on an example left hemisphere wrinkled (pial) cortical surface. **(B)** Example local gyrification index map in the same hemisphere. Higher values indicate areas of higher folding (more sulcation; see scale) (37). Abbreviations are as follows: anterior cingulate cortex (ACC) caudal ACC (cACC), rostral ACC (ACC), superior frontal cortex (SFC), local gyrification index (LGI), left hemisphere (lh), right hemisphere (rh).

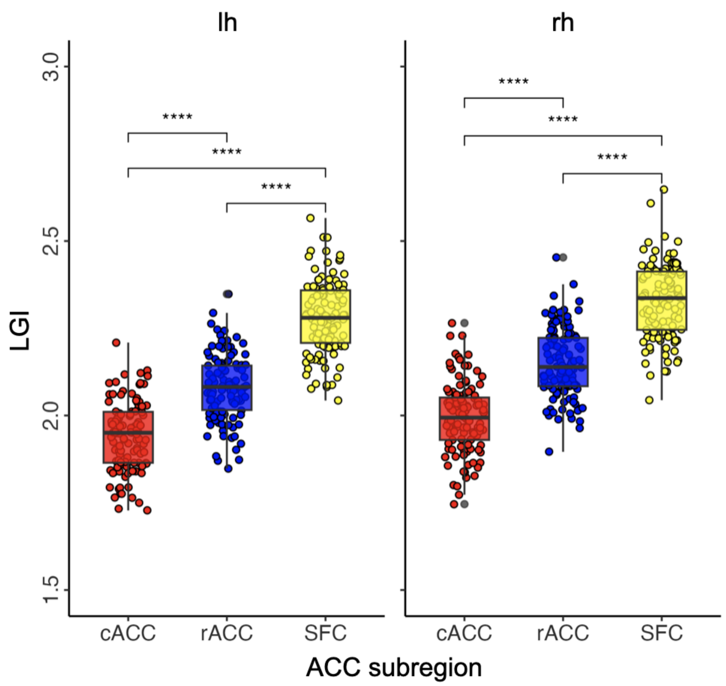


**Figure S3. Mean LGI values of ACC subregions.** Boxplot and scatterplot showing mean LGI values (y-axis) as a function of ACC subregion (Figure S2A; x-axis) and hemisphere (left and right plots). The significance of the effect of ACC subregion on the mean LGI (via t-tests) is shown with asterisks. Note that LGI values were also higher in the rh than lh (t-test, $p < .0001$). Abbreviations are as follows: anterior cingulate cortex (ACC) caudal ACC (cACC), rostral ACC (ACC), superior frontal cortex (SFC), local gyrification index (LGI), left hemisphere (lh), right hemisphere (rh). ****$p < .0001$

### Supplementary Tables

#### Table S1

*ANOVAs Regressing Impulsivity on Age and Gender*

Model 1: Predicting Pervasive Influence of Feelings

| Metric | F-value (df,df) | *p*-value | $\eta_p^2$ |
|---|---|---|---|
| Age | 3.22 (1,116) | .15 | 0.03 |
| Gender | 0.73 (2,116) | .48 | 0.01 |

Model 2: Predicting Feelings Trigger Action

| Metric | F-value (df,df) | *p*-value | $\eta_p^2$ |
|---|---|---|---|
| Age | 0.98 (1,116) | .65 | < 0.01 |
| Gender | 0.10 (2,116) | .90 | < 0.01 |

Model 3: Predicting Lack of Follow Through

| Metric | F-value (df,df) | *p*-value | $\eta_p^2$ |
|---|---|---|---|
| Age | 0.46 (1,116) | .71 | < 0.01 |
| Gender | 0.34 (2,116) | .71 | < 0.01 |

*Note.* Abbreviations are as follows: analysis of variance (ANOVA); degrees of freedom (df); partial eta-squared ($\eta_p^2$).

**Table S2**

*ANOVAs Regressing PCGS Length on Age and Gender*

| Model 1: Left PCGS Length (mm) | | | |
|---|---|---|---|
| Metric | F-value (df,df) | *p*-value | $\eta_p^2$ |
| Age | 0.15 (1,86) | .79 | < 0.01 |
| Gender | 0.23 (2,86) | .79 | < 0.01 |

| Model 2: Right PCGS Length (mm) | | | |
|---|---|---|---|
| Metric | F-value (df,df) | *p*-value | $\eta_p^2$ |
| Age | 0.35 (1,64) | .79 | < 0.01 |
| Gender | 0.23 (2,64) | .79 | < 0.01 |

| Model 3: PCGS Laterality Ratio (mm) | | | |
|---|---|---|---|
| Metric | F-value (df,df) | *p*-value | $\eta_p^2$ |
| Age | 0.03 (1,116) | .85 | < 0.01 |
| Gender | 0.40 (2,116) | .85 | < 0.01 |

*Note.* Abbreviations are as follows: analysis of variance (ANOVA); degrees of freedom (df); paracingulate sulcus (PCGS); partial eta-squared ($\eta_p^2$).

**Table S3**

*ANOVAs Regressing ACC LGI on Age and Gender*

Model 1: Right Caudal ACC

| Metric | F-value (df,df) | *p*-value | $\eta_p^2$ |
|---|---|---|---|
| Age | 15.54 (1, 109) | .0002 | 0.13 |
| Gender | 2.90 (2,109) | .059 | 0.05 |

Model 2: Left Caudal ACC

| Metric | F-value (df,df) | *p*-value | $\eta_p^2$ |
|---|---|---|---|
| Age | 9.49 (1,109) | .006 | 0.08 |
| Gender | 4.11 | .019 | 0.07 |

Model 3: Right Superior Frontal

| Metric | F-value (df,df) | *p*-value | $\eta_p^2$ |
|---|---|---|---|
| Age | 25.90 (1,109) | .000002 | 0.19 |
| Gender | 3.01 | .053 | .05 |

Model 4: Left Superior Frontal

| Metric | F-value (df,df) | *p*-value | $\eta_p^2$ |
|---|---|---|---|
| Age | 21.90 (1,109) | .00001 | 0.17 |
| Gender | 3.49 (2,109) | .034 | 0.06 |

Model 5: Right Rostral ACC

| Metric | F-value (df,df) | *p*-value | $\eta_p^2$ |
|---|---|---|---|
| Age | 12.45 (1,109) | .0002 | 0.10 |

| | | | |
|---|---|---|---|
| Gender | 2.13 (2,109) | .12 | 0.04 |

Model 6: Left Rostral ACC

| Metric | F-value (df,df) | *p*-value | $\eta_p^2$ |
|---|---|---|---|
| Age | 12.45 (1,109) | .001 | 0.10 |
| Gender | 1.17 (2,109) | .31 | 0.02 |

Model 7: Superior Frontal Laterality Ratio

| Metric | F-value (df,df) | *p*-value | $\eta_p^2$ |
|---|---|---|---|
| Age | 1.47 (1,109) | .45 | 0.01 |
| Gender | 0.33 (2,1090 | .71 | < 0.01 |

Model 8: Rostral ACC Laterality Ratio

| Metric | F-value (df,df) | *p*-value | $\eta_p^2$ |
|---|---|---|---|
| Age | 0.36 (1,109) | .54 | < 0.01 |
| Gender | 0.71 (2,109) | .54 | 0.01 |

Model 9: Rostral ACC Laterality Ratio

| Metric | F-value (df,df) | *p*-value | $\eta_p^2$ |
|---|---|---|---|
| Age | 0.34 (1,109) | .66 | < 0.01 |
| Gender | 0.42 (2,109) | .66 | < 0.01 |

*Note.* Abbreviations are as follows: anterior cingulate cortex (ACC); analysis of variance (ANOVA); degrees of freedom (df); partial eta-squared ($\eta_p^2$).

**Table S4**

*Multivariate Models Regressing Impulsivity on PCGS Length (mm)*

Model 1: Predicting Pervasive Influence of Feelings

| Metric | β-value | Std. Err. | *t*-value | *p*-value |
|---|---|---|---|---|
| Left PCGS Length | 0.002 | 0.004 | 0.52 | .78 |
| Right PCGS Length | 0.001 | 0.005 | 0.20 | .83 |

Multiple $R^2 < 0.01$, Adjusted $R^2 < 0.01$, $F(2,49) = 0.15$, $p = .86$

Model 2: Predicting Feelings Trigger Action

| Metric | β-value | Std. Err. | *t*-value | *p*-value |
|---|---|---|---|---|
| Left PCGS Length | 0.003 | 0.005 | 0.61 | .78 |
| Right PCGS Length | 0.002 | 0.005 | 0.35 | .82 |

Multiple $R^2 < 0.01$, Adjusted $R^2 < 0.01$, $F(2,49) = 0.24$, $p = .79$

Model 3: Predicting Lack of Follow Through

| Metric | β-value | Std. Err. | *t*-value | *p*-value |
|---|---|---|---|---|
| Left PCGS Length | 0.002 | 0.004 | 0.57 | .78 |
| Right PCGS Length | 0.005 | 0.004 | 1.05 | .78 |

Multiple $R^2 = 0.03$, Adjusted $R^2 < 0.01$, $F(2,49) = 0.69$, $p = .50$

*Note.* Abbreviations are as follows: paracingulate sulcus (PCGS); standard error (Std. Err.).

**Table S5**

*Multivariate Models Regressing Impulsivity on PCGS Laterality Ratio (mm)*

Model 1: Predicting Pervasive Influence of Feelings

| Metric | β-value | Std. Err. | *t*-value | *p*-value |
|---|---|---|---|---|
| PCGS Laterality Ratio | -0.13 | 0.10 | -1.30 | .78 |

Multiple $R^2 = 0.01$, Adjusted $R^2 < 0.01$, $F(1,118) = 1.69$, $p = .78$

Model 2: Predicting Feelings Trigger Action

| Metric | β-value | Std. Err. | *t*-value | *p*-value |
|---|---|---|---|---|
| Left PCGS Length | -0.08 | 0.10 | -0.81 | .78 |

Multiple $R^2 < 0.01$, Adjusted $R^2 < 0.01$, $F(1,118) = 0.66$, $p = .78$

Model 3: Predicting Lack of Follow Through

| Metric | β-value | Std. Err. | *t*-value | *p*-value |
|---|---|---|---|---|
| Left PCGS Length | -0.27 | 0.10 | -2.54 | .11 |

Multiple $R^2 = 0.05$, Adjusted $R^2 = 0.04$, $F(1,118) = 6.45$, $p = .11$

*Note.* Abbreviations are as follows: paracingulate sulcus (PCGS); standard error (Std. Err.).

**Table S6**

*Multivariate Models Regressing Impulsivity on ACC LGI*

Model 1: Predicting Pervasive Influence of Feelings

| Metric | β-value | Std. Err. | *t*-value | *p*-value |
|---|---|---|---|---|
| Right Caudal ACC | 1.34 | 1.51 | 0.88 | .87 |
| Left Caudal ACC | -0.56 | 1.60 | -0.35 | .87 |
| Right Superior Frontal | 0.23 | 1.50 | 0.15 | .87 |
| Left Superior Frontal | 0.41 | 1.49 | 0.28 | .87 |
| Right Rostral ACC | -0.23 | 1.48 | -0.16 | .87 |
| Left Rostral ACC | 0.82 | 1.50 | 0.55 | .87 |

Multiple $R^2$ = 0.05, Adjusted $R^2$ < 0.01, F(6,106) = 0.91, *p* = .49

Model 2: Predicting Feelings Trigger Action

| Metric | β-value | Std. Err. | *t*-value | *p*-value |
|---|---|---|---|---|
| Right Caudal ACC | 1.54 | 1.43 | 1.08 | .66 |
| Left Caudal ACC | 0.93 | 1.52 | 0.61 | .75 |
| Right Superior Frontal | 0.49 | 1.43 | 0.34 | .84 |
| Left Superior Frontal | 0.28 | 1.41 | 0.20 | .84 |
| Right Rostral ACC | -2.37 | 1.40 | -1.69 | .32 |
| Left Rostral ACC | -0.91 | 1.42 | -0.64 | .75 |

Multiple $R^2$ = 0.04, Adjusted $R^2$ < 0.01, F(6,106) = 0.237, *p* = .79

Model 3: Predicting Lack of Follow Through

| Metric | β-value | Std. Err. | *t*-value | *p*-value |
|---|---|---|---|---|
| Right Caudal ACC | 0.58 | 1.55 | 1.80 | .27 |

| | | | | |
|---|---|---|---|---|
| Left Caudal ACC | 0.03 | 1.65 | 0.02 | .98 |
| Right Superior Frontal | 2.75 | 1.55 | 1.78 | .27 |
| Left Superior Frontal | -1.25 | 1.54 | -0.82 | .72 |
| Right Rostral ACC | -0.62 | 1.52 | -0.41 | .82 |
| Left Rostral ACC | -1.73 | 1.54 | -1.12 | .61 |

Multiple $R^2$ = 0.05, Adjusted $R^2$ < 0.01, F(6,106) = 0.93, *p* = .47

*Note.* Abbreviations are as follows: anterior cingulate cortex (ACC); local gyrification index (LGI); standard error (Std. Err.).

**Table S7**

*Multivariate Models Regressing Impulsivity on ACC LGI Laterality Ratios*

Model 1: Predicting Pervasive Influence of Feelings

| Metric | β-value | Std. Err. | *t*-value | *p*-value |
|---|---|---|---|---|
| Caudal ACC | 3.22 | 5.38 | 0.60 | .93 |
| Superior Frontal | 0.49 | 6.45 | 0.08 | .93 |
| Rostral ACC | -1.40 | 5.40 | -0.26 | .93 |

Multiple $R^2 < 0.01$, Adjusted $R^2 < 0.01$, $F(3,109) = 0.17$, $p = .91$

Model 2: Predicting Feelings Trigger Action

| Metric | β-value | Std. Err. | *t*-value | *p*-value |
|---|---|---|---|---|
| Caudal ACC | 1.46 | 5.08 | 0.29 | .83 |
| Superior Frontal | -1.28 | 6.08 | -0.21 | .83 |
| Rostral ACC | -2.85 | 5.10 | -0.56 | .83 |

Multiple $R^2 < 0.01$, Adjusted $R^2 < 0.01$, $F(3,109) = 0.12$, $p = .94$

Model 3: Predicting Lack of Follow Through

| Metric | β-value | Std. Err. | *t*-value | *p*-value |
|---|---|---|---|---|
| Caudal ACC | 0.52 | 5.46 | 0.10 | .92 |
| Superior Frontal | 8.15 | 6.55 | 1.25 | .43 |
| Rostral ACC | 3.17 | 5.48 | 0.58 | .72 |

Multiple $R^2 = 0.02$, Adjusted $R^2 < 0.01$, $F(3,109) = 0.93$, $p = .47$

*Note.* Abbreviations are as follows: anterior cingulate cortex (ACC); local gyrification index (LGI); standard error (Std. Err.).

**Table S8**

*χ2 Tests Relating PCGS Presence or Asymmetry to SCID-5 MDD Lifetime Diagnosis*

| Left Hemisphere PCGS Presence | MDD | | Analysis |
|---|---|---|---|
| | Yes | No | |
| Present | 73 (79.3%) | 19 (20.7%) | $\chi 2(1) = 0.23$, $p = .63$ |
| Absent | 24 (85.7%) | 4 (14.3%) | |

| Right Hemisphere PCGS Presence | MDD | | Analysis |
|---|---|---|---|
| | Yes | No | |
| Present | 56 (82.4%) | 12 (17.6%) | $\chi 2(1) = 0.06$, $p = .80$ |
| Absent | 41 (78.8%) | 11 (21.2%) | |

| PCGS Asymmetry | MDD | | Analysis |
|---|---|---|---|
| | Yes | No | |
| Symmetric | 56 (84.8%) | 10 (15.2%) | $\chi 2(2) = 1.75$, $p = .41$ |
| LW Asymmetric | 29 (74.4%) | 10 (25.6%) | |
| RW Asymmetric | 12 (80%) | 3 (20%) | |

*Note.* Incidence rates are presented as n (%). Abbreviations are as follows: chi-squared ($\chi 2$); degrees of freedom (df); leftward (LW); major depressive disorder (MDD); paracingulate sulcus (PCGS); rightward (RW).

**Table S9**

*χ2 Tests Relating PCGS Presence or Asymmetry to SCID-5 AD Lifetime Diagnosis*

| Left Hemisphere PCGS Presence | AD | | Analysis |
|---|---|---|---|
| | Yes | No | |
| Present | 60 (65.2%) | 32 (34.8%) | $\chi 2(1) = 0.54$, $p = .46$ |
| Absent | 21 (75%) | 7 (25%) | |

| Right Hemisphere PCGS Presence | AD | | Analysis |
|---|---|---|---|
| | Yes | No | |
| Present | 43 (63.2%) | 25 (36.8%) | $\chi 2(1) = 0.89$, $p = .34$ |
| Absent | 38 (73.1%) | 14 (26.9%) | |

| PCGS Asymmetry | AD | | Analysis |
|---|---|---|---|
| | Yes | No | |
| Symmetric | 40 (60.6%) | 26 (39.4%) | $\chi 2(2) = 3.33$, $p = .18$ |
| LW Asymmetric | 29 (74.4%) | 10 (25.6%) | |
| RW Asymmetric | 12 (80%) | 3 (20%) | |

*Note.* Incidence rates are presented as n (%). Abbreviations are as follows: anxiety disorder (AD); chi-squared (*χ2*); degrees of freedom (df); leftward (LW); paracingulate sulcus (PCGS); rightward (RW).

**Table S10**

*χ2 Tests Relating PCGS Presence or Asymmetry to SCID-5 AUD Lifetime Diagnosis*

| Left Hemisphere PCGS Presence | *AUD* | | Analysis |
|---|---|---|---|
| | Yes | No | |
| Present | 21 (22.8%) | 71 (77.2%) | *χ2*(1) = 0.50, *p* = .47 |
| Absent | 4 (14.3%) | 24 (85.7%) | |

| Right Hemisphere PCGS Presence | AUD | | Analysis |
|---|---|---|---|
| | Yes | No | |
| Present | 13 (19.1%) | 55 (90.9%) | *χ2*(1) = 0.09, *p* = .76 |
| Absent | 12 (23.1%) | 40 (76.9%) | |

| PCGS Asymmetry | AUD | | Analysis |
|---|---|---|---|
| | Yes | No | |
| Symmetric | 13 (19.7%) | 53 (80.3%) | *χ2*(2) = 1.11, *p* = .57 |
| LW Asymmetric | 10 (25.6%) | 29 (74.4%) | |
| RW Asymmetric | 2 (13.3%) | 13 (86.7%) | |

*Note.* Incidence rates are presented as n (%). Abbreviations are as follows: alcohol use disorder (AUD); chi-squared (*χ2*); degrees of freedom (df); leftward (LW); paracingulate sulcus (PCGS); rightward (RW).

**Table S11**

*χ2 Tests Relating PCGS Presence or Asymmetry to SCID-5 SUD Lifetime Diagnosis*

| Left Hemisphere PCGS Presence | SUD | | Analysis |
|---|---|---|---|
| | Yes | No | |
| Present | 21 (22.8%) | 71 (77.2%) | $\chi 2(1) = 1.28$, $p = .25$ |
| Absent | 3 (10.7%) | 25 (89.3%) | |

| Right Hemisphere PCGS Presence | SUD | | Analysis |
|---|---|---|---|
| | Yes | No | |
| Present | 15 (22.1%) | 53 (77.9%) | $\chi 2(1) = 0.17$, $p = .67$ |
| Absent | 9 (17.3%) | 43 (82.7%) | |

| PCGS Asymmetry | SUD | | Analysis |
|---|---|---|---|
| | Yes | No | |
| Symmetric | 14 (21.2%) | 52 (77.8%) | $\chi 2(2) = 0.48$, $p = .78$ |
| LW Asymmetric | 8 (20.5%) | 31 (79.5%) | |
| RW Asymmetric | 2 (13.3%) | 13 (86.7%) | |

*Note.* Incidence rates are presented as n (%). Abbreviations are as follows: chi-squared (*χ2*); degrees of freedom (df); leftward (LW); paracingulate sulcus (PCGS); rightward (RW); substance use disorder (SUD).

**Table S12**

*ANOVAs Regressing Age on PCGS Presence and Asymmetry*

| Metric | F-value (df,df) | *p*-value | $\eta_p^2$ |
|---|---|---|---|
| Left PCGS Presence | 0.28 (1,117) | .89 | < 0.01 |
| Right PCGS Presence | 0.39 (1,117) | .89 | < 0.01 |
| PCGS Asymmetry | 0.04 (2,117) | .96 | < 0.01 |

*Note.* Abbreviations are as follows: analysis of variance (ANOVA); degrees of freedom (df); paracingulate sulcus (PCGS); partial eta-squared ($\eta_p^2$).

**Table S13**

*χ2 Tests Relating PCGS Presence and Asymmetry to Gender*

| Left Hemisphere PCGS Presence | Gender | | | Analysis |
|---|---|---|---|---|
| | M | F | NB | |
| Present | 29 (31.5%) | 59 (64.1%) | 4 (4.4%) | $\chi2(2) = 0.51$, $p = .77$ |
| Absent | 7 (25%) | 20 (71.4%) | 1 (3.6%) | |

| Right Hemisphere PCGS Presence | Gender | | | Analysis |
|---|---|---|---|---|
| | M | F | NB | |
| Present | 24 (35.3%) | 42 (61.8%) | 2 (2.9%) | $\chi2(2) = 2.43$, $p = .29$ |
| Absent | 12 (23.1%) | 37 (71.1%) | 3 (5.8%) | |

| PCGS Asymmetry | Gender | | | Analysis |
|---|---|---|---|---|
| | M | F | NB | |
| Symmetric | 21 (31.8%) | 42 (63.6%) | 3 (4.6%) | $\chi2(4) = 1.23$, $p = .87$ |
| LW Asymmetric | 10 (25.6%) | 27 (69.2%) | 2 (5.2%) | |
| RW Asymmetric | 5 (33.3%) | 10 (66.7%) | 0 (0%) | |

*Note.* Incidence rates are presented as n (%). Abbreviations are as follows: chi-squared ($\chi2$); degrees of freedom (df); female (F); male (M); leftward (LW); major depressive disorder (MDD); nonbinary (NB); paracingulate sulcus (PCGS); rightward (RW).

**Table S14**

*ANOVAs Regressing Impulsivity on PCGS Presence*

Model 1: Predicting Pervasive Influence of Feelings

| Metric | F-value (df,df) | *p*-value | $\eta_p^2$ |
|---|---|---|---|
| Left PCGS | 2.13 (1,117) | .29 | 0.02 |
| Right PCGS | >0.01 (1,117) | .99 | < 0.01 |

Model 2: Predicting Feelings Trigger Action

| Metric | F-value (df,df) | *p*-value | $\eta_p^2$ |
|---|---|---|---|
| Left PCGS | 1.21 (1,117) | .29 | 0.01 |
| Right PCGS | 0.08 (1,117) | .99 | < 0.01 |

Model 3: Predicting Lack of Follow Through

| Metric | F-value (df,df) | *p*-value | $\eta_p^2$ |
|---|---|---|---|
| Left PCGS | 23.87 (1,117) | .00001 | 0.17 |
| Right PCGS | >0.01 (1,117) | .99 | < 0.01 |

*Note.* Abbreviations are as follows: analysis of variance (ANOVA); degrees of freedom (df); paracingulate sulcus (PCGS); partial eta-squared ($\eta_p^2$).

**Table S15**

*ANOVAs Regressing Impulsivity on PCGS Asymmetry*

Model 1: Predicting Pervasive Influence of Feelings

| Metric | F-value (df,df) | *p*-value | $\eta_p^2$ |
|---|---|---|---|
| PCGS Asymmetry | 0.79 (2,117) | .65 | 0.01 |

Model 2: Predicting Feelings Trigger Action

| Metric | F-value (df,df) | *p*-value | $\eta_p^2$ |
|---|---|---|---|
| PCGS Asymmetry | 0.43 (2,117) | .65 | <0.01 |

Model 3: Predicting Lack of Follow Through

| Metric | F-value (df,df) | *p*-value | $\eta_p^2$ |
|---|---|---|---|
| PCGS Asymmetry | 12.69 (2,117) | .00003 | 0.18 |

*Note.* Abbreviations are as follows: analysis of variance (ANOVA); degrees of freedom (df); paracingulate sulcus (PCGS); partial eta-squared ($\eta_p^2$).

Prenatal Gyrification Pattern Affects Age at Onset in Frontotemporal Dementia. *Cereb Cortex*. https://doi.org/10.1093/cercor/bhab457

26. Ramos Benitez J, Kannan S, Hastings WL, Parker BJ, Willbrand EH, Weiner KS (2024): Ventral temporal and posteromedial sulcal morphology in autism spectrum disorder.

*Neuropsychologia* 195: 108786.

37. Schaer M, Cuadra MB, Schmansky N, Fischl B, Thiran J-P, Eliez S (2012): How to measure cortical folding from MR images: a step-by-step tutorial to compute local gyrification index. *J Vis Exp* e3417.
